# IFT25 is required for the construction of the trypanosome flagellum

**DOI:** 10.1101/479642

**Authors:** Diego Huet, Thierry Blisnick, Sylvie Perrot, Philippe Bastin

## Abstract

Intraflagellar transport (IFT), the movement of protein complexes responsible for the assembly of cilia and flagella, is remarkably well conserved from protists to humans. However, two IFT components (IFT25 and IFT27) are missing from multiple unrelated eukaryotic species. In mouse, IFT25 and IFT27 are not required for assembly of several cilia with the noticeable exception of the flagellum of spermatozoa. Here we show that the *Trypanosoma brucei* IFT25 protein is a proper component of the IFT-B complex and displays typical IFT trafficking. Using bimolecular fluorescence complementation assays, we reveal that IFT25 and IFT27 interact within the flagellum in live cells during the IFT transport process. IFT25-depleted cells construct tiny disorganised flagella that accumulate IFT-B proteins (with the exception of IFT27, the binding partner of IFT25) but not IFT-A proteins. This phenotype is comparable to the one following depletion of IFT27 and shows that IFT25/IFT27 constitute a specific module requested for proper IFT and flagellum construction in trypanosomes. We discuss the possible reasons why IFT25/IFT27 would be required for only some types of cilia.

## Introduction

Cilia and flagella are organelles conserved across eukaryotes and perform variable functions ranging from motility to sensing or morphogenesis. Intraflagellar transport (IFT) is the movement of protein particles or trains that delivers tubulin and other precursors at the tip of cilia and flagella for elongation (Craft et al., 2015; Kozminski et al., 1993). Trains are moved by the action of specific molecular motors of the kinesin and dynein family during anterograde and retrograde transport, respectively. They are composed of polymers made of two protein complexes termed IFT-A (6 proteins) and IFT-B (16 proteins) (Cole et al., 1998; Piperno and Mead, 1997; Taschner and Lorentzen, 2016). *IFT* genes are evolutionary conserved, being found in most eukaryotic organisms with cilia and flagella ranging from protists to mammals (van Dam et al., 2013). The only major exceptions are encountered in *Plasmodium* subspecies that assemble their flagella in the cytoplasm and whose genomes lack all *IFT* genes (Briggs et al., 2004; Sinden et al., 1976). Inhibition of IFT blocks the construction of cilia and flagella in all species investigated to date (Brown et al., 1999; Han et al., 2003; Kohl et al., 2003; Kozminski et al., 1995; Marszalek et al., 2000). In general, absence of an IFT-B protein results in failure to construct the organelle whereas absence of an IFT-A protein interferes with retrograde transport, leading to the accumulation of IFT trains and to the formation of shorter cilia (Absalon et al., 2008b; Blacque et al., 2006; Efimenko et al., 2006; Iomini et al., 2009).

A noticeable exception to this conservation is IFT25 that is missing from the genome of 15 ciliated species (van Dam et al., 2013)(Table 1). IFT25 was first identified as a candidate chaperone protein based on some sequence homology with heat shock proteins (Bellyei et al., 2007). However, this was not supported by structural analysis of the *Chlamydomonas* IFT25 protein (Bhogaraju et al., 2011). IFT25 was found to be a member of the IFT-B complex in both *Chlamydomonas* and human cells (Follit et al., 2009; Lechtreck et al., 2009b). It associates with the small G protein IFT27 (also known as RABL4) and is added to the IFT-B complex just prior to IFT activation (Wang et al., 2009). The crystal structure of the IFT25/IFT27 dimer was determined and revealed tight interaction between the two proteins (Bhogaraju et al., 2011). IFT25 and IFT27 participate to the B1 sub-complex of the IFT-B together with 7 other IFT-B proteins (Taschner et al., 2014; Taschner et al., 2016). In species missing IFT25, IFT27 is often (but not always) absent (Table 1).

**Table 1.** List of species missing IFT25 and/or IFT27

Data from (van Dam et al., 2013)
| Missing gene | Species |
| --- | --- |
| IFT25 | *Monosiga, Thalassosira, Emiliana, Paramecium, Tetrahymena* |
| IFT25+IFT27 | *Drosophila, Giardia, Ciona, Anopheles, C. elegans, Physcomitrella, Selaginella* |
| IFT27 | none |

The loss of IFT25 in multiple different species raises the question of its actual role in cilia construction and function. In contrast to other *IFT* genes, *IFT25* gene deletion in mouse was viable until birth. Cilia in the lung and in the trachea were present and exhibited a normal morphology, suggesting that the protein was not essential for the assembly of cilia. Embryonic fibroblasts of the *ift25* mutant contained all examined IFT proteins with the exception of IFT27 (Keady et al., 2012). Nevertheless, embryo deprived of IFT25 died at birth with multiple defects related to dysfunctions in hedgehog signalling. Detailed investigation confirmed that the distribution of protein components of this key signalling pathway was drastically modified both in quiescent and active situations, leading to a model where IFT25 (Keady et al., 2012) and IFT27 (Eguether et al., 2014; Liew et al., 2014) would contribute to the transport of signalling molecules. This could be mediated by the BBSome, another protein complex initially identified because of its association to the Bardet-Biedl Syndrome (Eguether et al., 2014; Liew et al., 2014). However, specific deletion of IFT25 and IFT27 in male germ cells revealed that both proteins are essential for the assembly of the flagellum in spermatozoa (Liu et al., 2017; Zhang et al., 2017).

Intriguingly, IFT25 is conserved in *Chlamydomonas*, *Volvox* and several protists (*Trichomonas, Trypanosoma*) where the hedgehog cascade is absent, raising the question of its role in these organisms. Inhibition of IFT25 expression in *Chlamydomonas* did not visibly affect neither IFT nor flagellum construction but perturbed BBSome trafficking (Dong et al., 2017). In an effort to clarify the function of IFT25, we investigated its function in the protist *Trypanosoma brucei*. This organism is well known for being responsible for sleeping sickness in Africa but also turned out to be an excellent model to study flagellum construction (Vincensini et al., 2011). It possesses a single flagellum that is replicated at each cell cycle whilst maintaining the existing one, providing the opportunity to compare growing and mature flagella in the same cell (Sherwin and Gull, 1989). All 22 *IFT* genes are conserved in *T. brucei*, flagellar trafficking was demonstrated upon expression of various fusions with fluorescent proteins and inhibition of *IFT* gene expression by RNAi knockdown severely impedes flagellum construction (Absalon et al., 2008b; Adhiambo et al., 2009; Blisnick et al., 2014; Buisson et al., 2013; Davidge et al., 2006; Kohl et al., 2003). Moreover, affinity purification using IFT56 identified most members of the IFT-B complex (Franklin and Ullu, 2010), with the exception of IFT20, IFT25 and IFT27.

Here, we show that IFT25 is a *bona fide* member of the IFT-B complex in trypanosomes and formally show that it undergoes IFT trafficking. Bimolecular fluorescence complementation assays using split YFP fragments fused to IFT25 and IFT27 revealed the typical IFT movement of fluorescent proteins within the flagellum, demonstrating interaction between the two proteins during IFT. RNAi knockdown blocks flagellum construction and results in inhibition of retrograde transport, a phenotype previously reported for IFT27 silencing in the same organism (Huet et al., 2014). These results reveal that IFT25 and IFT27 form a unique module that is essential for IFT trafficking and flagellum construction in trypanosomes, with several similarities with their role in assembly of the sperm flagellum. We discuss potential relationships between the flagellum of trypanosomes and mammalian spermatozoa.

## Results

### IFT25 associates to the IFT-B complex and traffics inside the trypanosome flagellum

In *T. brucei*, IFT25 is encoded by gene Tb927.11.13300 and was detected in purified intact flagella of both insect (Subota et al., 2014) and bloodstream (Oberholzer et al., 2011) stages but not in detergent salt-extracted flagellar fractions (Broadhead et al., 2006), indicative of an association to the flagellum matrix. To investigate the exact localisation of IFT25 in *T. brucei*, a GFP::IFT25 fusion was expressed. Western blot analyses using an anti-GFP antibody showed a single band migrating between the 37 and the 50 kDa markers, in agreement with the expected mass of the fusion protein (45 kDa)(Figure 1A). The anti-GFP antibody was then used in immunofluorescence assays (IFA) along with the anti-IFT27 antibody to assess if both proteins co-localise inside the flagellar compartment. In GFP::IFT25 expressing cells, the anti-GFP antibody stained spots all along the flagellum, with a brighter signal at the base of the organelle (Figure 1B). Co-staining with the anti-IFT27 antibody (Huet et al., 2014) revealed a similar signal that co-localized in most parts of the flagellum with the GFP-staining (Figure 1B).

**Fig. 1.**
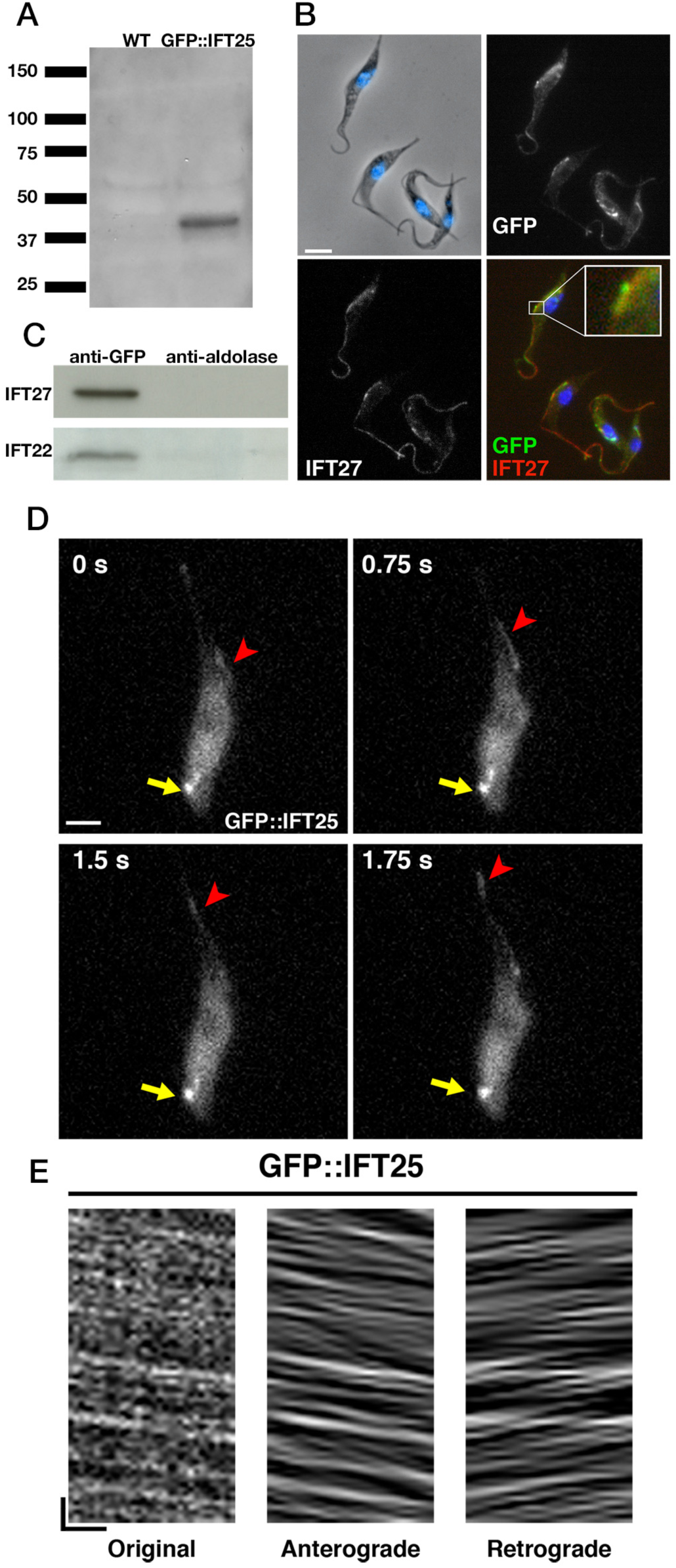
IFT25 is an IFT-B protein that displays intraflagellar transport. (A) Western blot analysis of total cell extracts from wild-type or GFP::IFT25 trypanosomes probed with an anti-GFP. A band of the expected electrophoretic motility is observed only in GFP::IFT25 cells. (B) Immunofluorescence analysis of cells expressing the GFP-tagged version of IFT25 fixed in methanol and stained with the anti-GFP antibody. The top left panel shows the phase-contrast image merged with DAPI (blue) staining, the top right one shows the anti-GFP staining (white) and the bottom left shows the anti-IFT27 staining (white). The bottom right panel shows the merged images (green for GFP and red for IFT27) and a close-up of the flagellum showing the partial colocalisation of the anti-GFP and anti-IFT27 signals. Scale bar: 5µm. (C) Immuno-precipitates of total proteins extracts from the cell line expressing GFP::IFT25 by an anti-GFP antibody or anti-aldolase antibody were separated on 4-15 % polyacrylamide gels. Corresponding immunoblots with the anti-IFT27 (top row) or the anti-IFT22 (bottom row) are shown. These results were replicated in two independent experiments. (D) Still images of a trypanosome expressing GFP::IFT25. The yellow arrow shows the fluorescent protein pool at the level of the flagellum base and the red arrows indicate the successive position of an anterograde IFT train at the indicated times. Scale bar: 2µm. (E) Kymograph from the same image sequence shows clear and typical IFT traces for GFP::IFT25. Anterograde and retrograde events were separated as previously described (Buisson et al., 2013). Horizontal scale bar is 2µm and vertical scale bar is 2 s.

IFT25 and IFT27 were not found in IFT-B complex purification, possibly because of their small size (Franklin and Ullu, 2010). To establish whether IFT25 was a member of the IFT-B complex, immuno-precipitation experiments were carried out on whole cell extracts from trypanosomes expressing GFP::IFT25. The anti-GFP was used for immuno-precipitation alongside an anti-aldolase as control and precipitates were analysed by western blotting using antibodies against either IFT22 (Adhiambo et al., 2009) or IFT27 (Huet et al., 2014), two recognised members of the IFT-B complex. The anti-GFP successfully immuno-precipitated IFT22 and IFT27, in contrast to the anti-aldolase (Fig. 1C). These results reveal that IFT25 is also a member of the IFT-B complex in trypanosomes.

To further link IFT25 to intraflagellar transport, live cells expressing GFP::IFT25 were observed by video microscopy. Transport in the anterograde direction was easily detected whereas retrograde transport was more discrete but clearly present upon careful examination (Video S1 and still images at Fig. 1D). Kymograph analyses (Fig. 1E) determined mean anterograde velocity to 2.7 ± 0.66 µm/s (n=229) while mean retrograde velocity was estimated to 3.1 ± 0.90 µm/s (n=296). Similar speed values have been observed for fluorescent fusion proteins with IFT52 (Buisson et al., 2013), IFT81 (Bertiaux et al., 2018; Bhogaraju et al., 2013) or IFT27 (Huet et al., 2014).

Since immuno-precipitations were done on whole cell extracts, we used bimolecular fluorescence complementation assays (Kerppola, 2008) to investigate whether IFT25 and IFT27 specifically interact. This approach had been used successfully to show interactions between IFT proteins or IFT proteins and transition fibre components at the base of cilia in *C. elegans* (Wei et al., 2013; Yi et al., 2017). In *Chlamydomonas*, it was proposed that IFT25 and IFT27 associate to the IFT-B complex at a late stage of assembly (Wang et al., 2009). To evaluate where does the interaction between the two proteins take place in *T. brucei*, we fused the N-terminal portion of YFP (aa1-155) to IFT25 and the shorter C-terminal portion (aa156-239) to IFT27 that contained a Ty-1 epitope (Bastin et al., 1996). Constructs were designed for endogenous tagging at the amino-terminal end of each protein, ensuring appropriate control of expression (Kelly et al., 2007). Expression of each protein individually did not lead to visible fluorescence in live cells but could be detected by IFA using a rabbit polyclonal antibody against GFP (Figure 2A-B). By contrast, fluorescent signals were easily detected in live cells when both proteins were co-expressed. Typical IFT movement was detected in all cells examined (Video S2, still images at Figure 2C). We conclude that IFT25 is a proper component of the IFT-B complex in *T. brucei* and demonstrate that the protein undergoes IFT within the flagellum while interacting with IFT27.

**Fig. 2.**
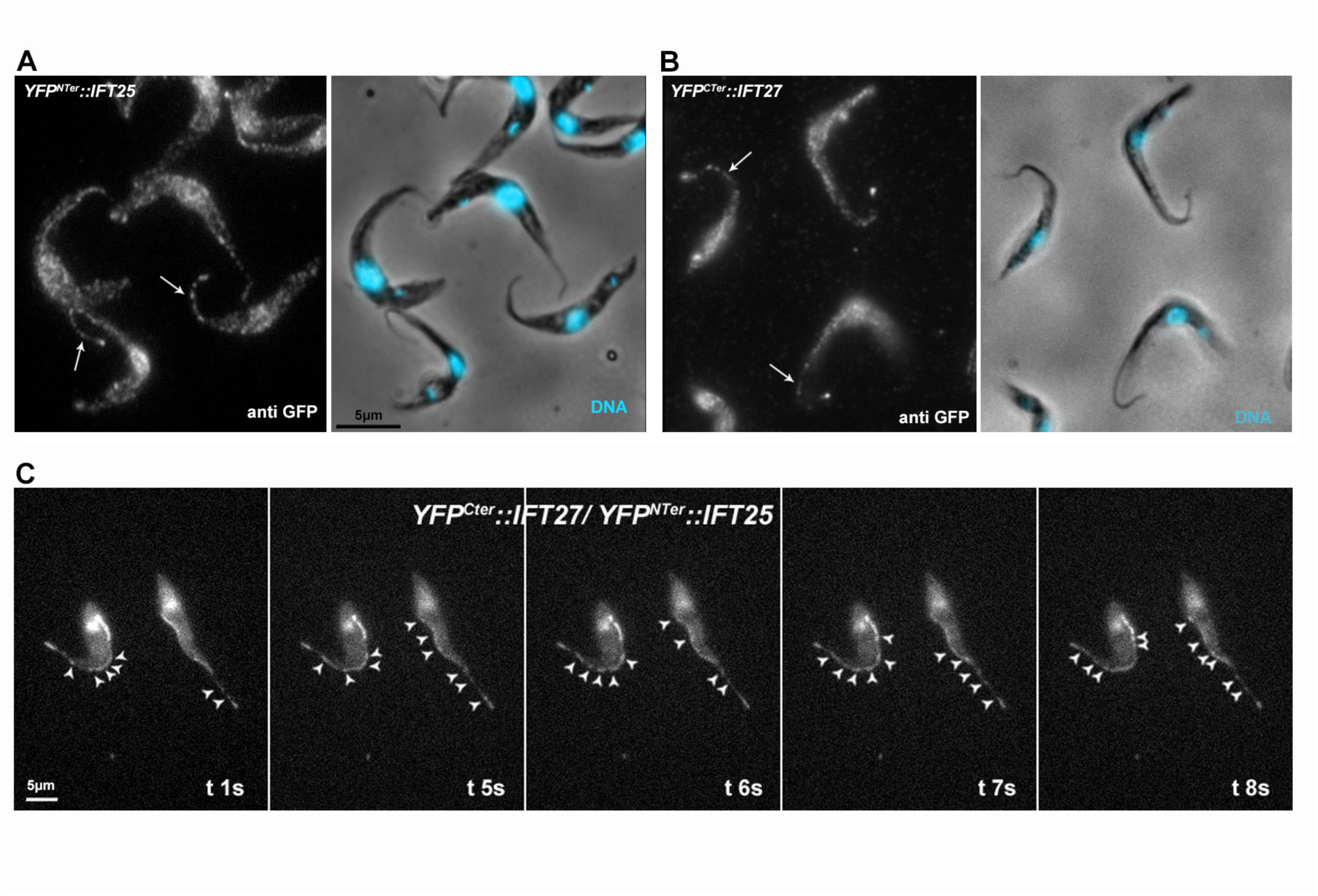
Bimolecular fluorescence complementation assay shows that IFT25 and IFT27 interact and traffic together during IFT. (A-B) Expression of the N-terminal part of YFP fused to IFT25 (A) or the C-terminal part of YFP fused to IFT27 (B) individually was detected by a polyclonal anti-GFP antiserum. The left panels show the immunofluorescence with the anti-GFP antiserum (white) and the right panels show the phase contrast image with DAPI signal to stain DNA-containing organelles (nucleus and kinetoplast). The arrows indicate flagella with a clear IFT signal pattern. (C) Simultaneous presence of IFT25 fused to the N-terminal part of YFP and of IFT27 fused to the C-terminal part of YFP results in reconstitution of YFP and emission of fluorescence. Still images at the indicating time points of live cells show trafficking of the interacting IFT25 and IFT27 proteins in the flagellum under the form of IFT trains (indicated with arrowheads). Video S2 shows the corresponding sequence of images.

### IFT25 is needed for flagellar assembly in *T. brucei*

Studies in mice showed that IFT25 is not needed for ciliary assembly in embryonic fibroblasts or in tracheal cells (Keady et al., 2012) but is essential for construction of the flagellum of spermatozoa (Liu et al., 2017; Zhang et al., 2017). To evaluate the role of IFT25 in *T. brucei*, an inducible RNA interference (RNAi) approach was used to deplete the expression of the protein in 29-13 procyclic trypanosomes in culture. The 29-13 strain expresses the tetracycline repressor and the viral T7 RNA polymerase, which enable tetracycline-dependent expression of double-stranded RNA leading to RNAi-mediated gene inhibition (Wang et al., 2000; Wirtz et al., 1999). We therefore generated the *IFT25^RNAi^* cell line and grew it in the presence of tetracycline to induce IFT25 knockdown. The first phenotype was an obvious growth defect upon depletion of the protein (Figure S1) and microscopy examination showed the presence of multiple immobile cells that were shorter than usual. Scanning electron microscopy analysis of non-induced cells showed the conventional morphology with elongated cells and a long flagellum (Figure 3A). In contrast, growth of *IFT25^RNAi^* cells in RNAi conditions for 24 (Figure 3B) or 48h (Figure 3C) revealed emergence of small cells with much shorter flagella. The flagellum is a central actor of cell morphogenesis in trypanosomes and contributes to the control of cell length (Kohl et al., 2003).

To examine more closely the nature of the flagellar defects, induced and non-induced *IFT25^RNAi^* cells were fixed and prepared for transmission electron microscopy (TEM) analysis. Sections through flagella or their base exhibited the normal aspect in control non-induced *IFT25^RNAi^* cells with the presence of the basal body, the transition zone and the axoneme and paraflagellar rod (Figure 3D). By contrast, induced cells looked dramatically different (Figure 3E-G). Their flagella were very short and the microtubule doublets frequently looked disorganised whilst a large amount of amorphous material accumulated towards the distal end of the organelle. Small vesicles were frequently detected among the amorphous material. This observation is reminiscent of a defect in retrograde transport as reported previously in *T. brucei* upon silencing of the IFT-A component IFT140 (Absalon et al., 2008a) or of any component of the IFT dynein complex (Blisnick et al., 2014). A similar phenotype has been observed for IFT22 (RAB-like 5)(Adhiambo et al., 2009) and IFT27 (RAB-like 4)(Huet et al., 2014), the two small G proteins members of the IFT-B complex.

**Fig. 3.**
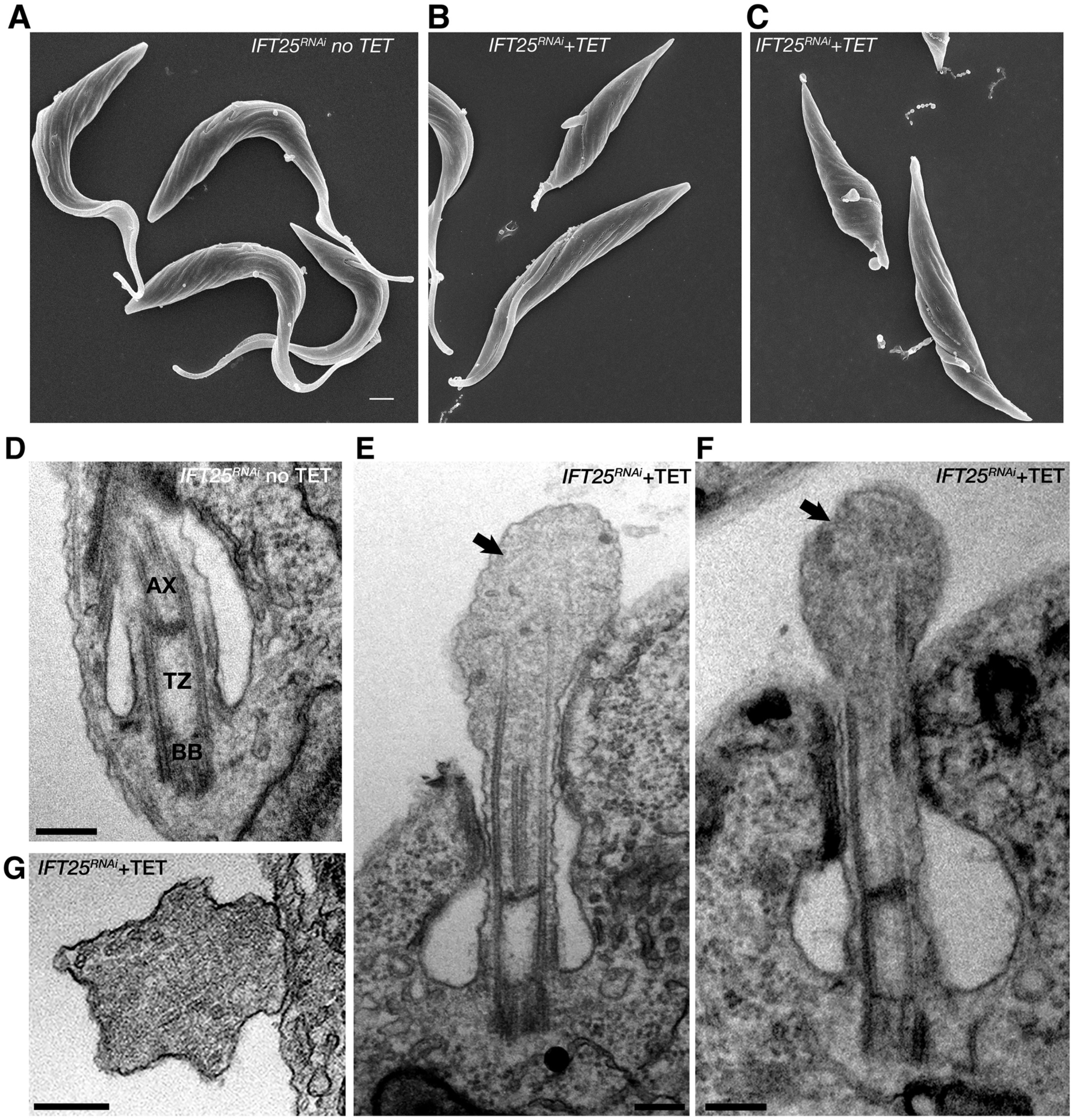
Knockdown of IFT25 inhibits flagellum construction. (A-C) Scanning electron microscopy images of *IFT25^RNAi^* cells either in non-induced conditions (A, no tet) or grown in the presence of tetracycline for 24 (B) or 48 hours (C). Scale bar: 1 µm. (D-G) Transmission electron microscopy images of sections from *IFT25^RNAi^* cells either in non-induced conditions (D, no tet) or grown in the presence of tetracycline for 48h (E-G). The presence of the basal body (BB), the transition zone (TZ) and the axoneme (AX) is easily identified in the non-induced control. Excessive IFT-like material is indicated with a black arrow. Scale bars: 200 nm.

Flagellum composition was analysed at the molecular level by IFA using simultaneously Mab25, a monoclonal antibody detecting an axoneme-specific protein (Dacheux et al., 2012) and a monoclonal antibody recognizing the IFT-B protein IFT172 (Absalon et al., 2008b). Both antibodies displayed the expected staining pattern in non-induced cells: a uniform distribution along the flagellum for Mab25 (typical of axonemal proteins), and a strong signal at the base of the flagellum and a succession of elongated spots along the flagellum for IFT172 (Figure 4A). By contrast, staining of *IFT25^RNAi^* cells where RNAi had been induced for two days revealed the presence of very short flagella with accumulation of excessive IFT172, further supporting a contribution of IFT25 to retrograde transport (Figure 4B). Taken together, these results show that IFT25 is required for flagellar assembly in *T. brucei* and that its depletion leads to an inhibition of retrograde IFT.

**Fig. 4.**
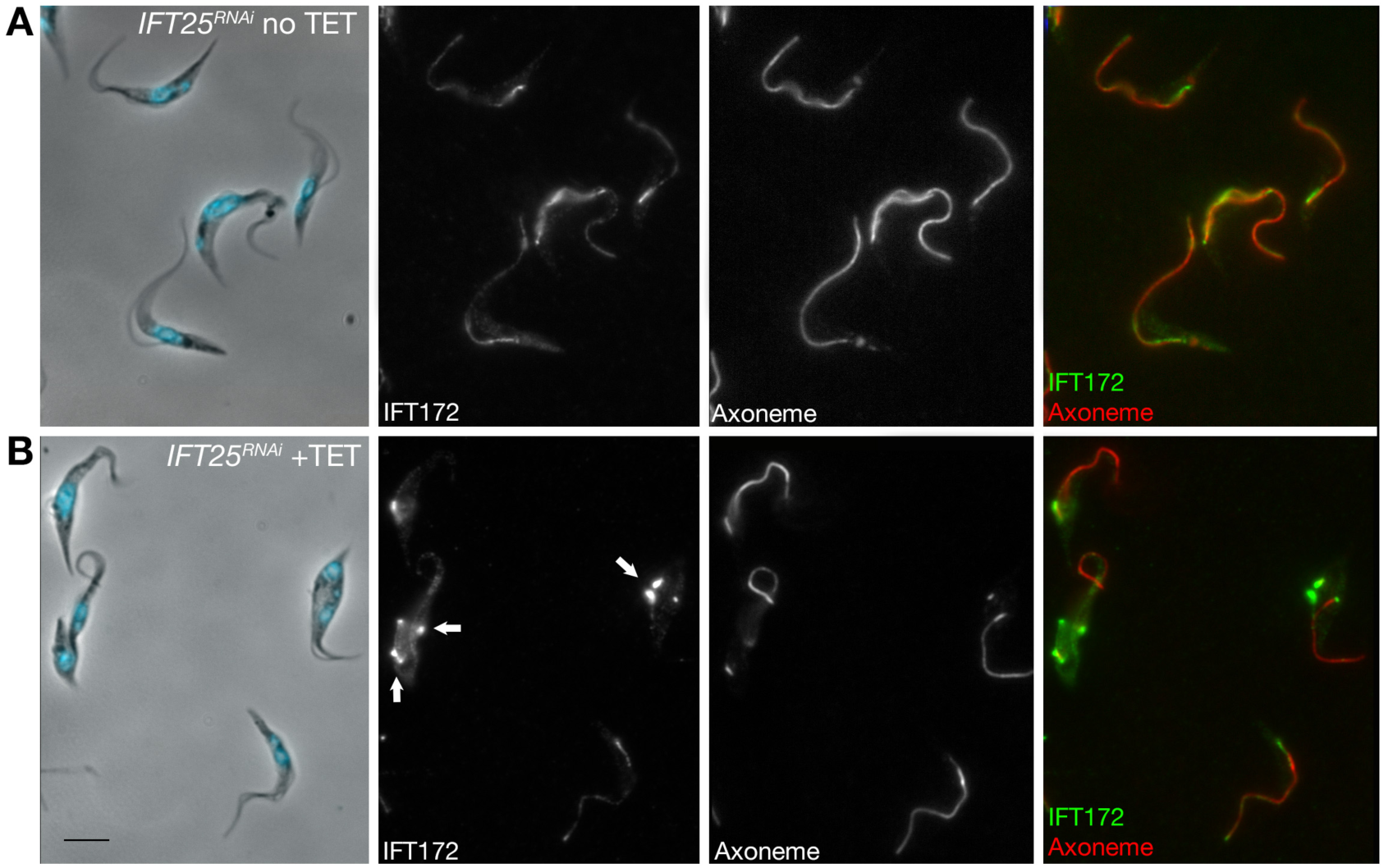
Knockdown of IFT25 results in accumulation of IFT-B proteins in the short flagellum, indicative of a retrograde transport phenotype. Non-induced (A) and 2-day induced (B) *IFT25^RNAi^* cells were fixed in methanol, stained with the mAb25 antibody (white, red on the merged panel) to detect the axoneme and the anti-IFT172 antibody (white, green on the merged panel) then counterstained with DAPI. The arrows show abnormally short axonemes with massive accumulation of IFT172 detected only in induced samples. Scale bar: ?µm.

### IFT25 is required for access to the flagellum of IFT27 and IFT140

We then analysed the fate of IFT proteins once the expression of IFT25 is inhibited in the *IFT25^RNAi^* cell line. Upon induction, no changes of the total steady-state amounts of IFT-B members IFT22, IFT27 or IFT172 were observed by western blot (Figure 5A). IFA using the anti-IFT27 antibody showed that the protein is found in the cytoplasm upon depletion of IFT25 and does not accumulate in short flagella in contrast to he classic IFT-B marker IFT172 (Figure 5B). To monitor the situation of the IFT-A complex, the *IFT25^RNAi^* cell line was transfected with a construct allowing expression of the IFT140 fused to a Tandem Tomato (TdT) marker from its endogenous locus (Huet et al., 2014). In non-induced conditions, TdT::IFT140 trafficked normally within the flagellum (Video S3, field view at Fig. 5C). In contrast, TdT::IFT140 was immotile in induced *IFT25^RNAi^* cells (Video S4) and appeared stuck at the base of the flagellum with little or no signal within the organelle (Figure 5D). Therefore, the absence of IFT25 prevents both IFT27 association to the IFT-B complex and IFT-A proteins access the flagellum. The absence of IFT-A could explain the retrograde phenotype exactly as observed for IFT27 knockdown (Huet et al., 2014).

**Fig. 5.**
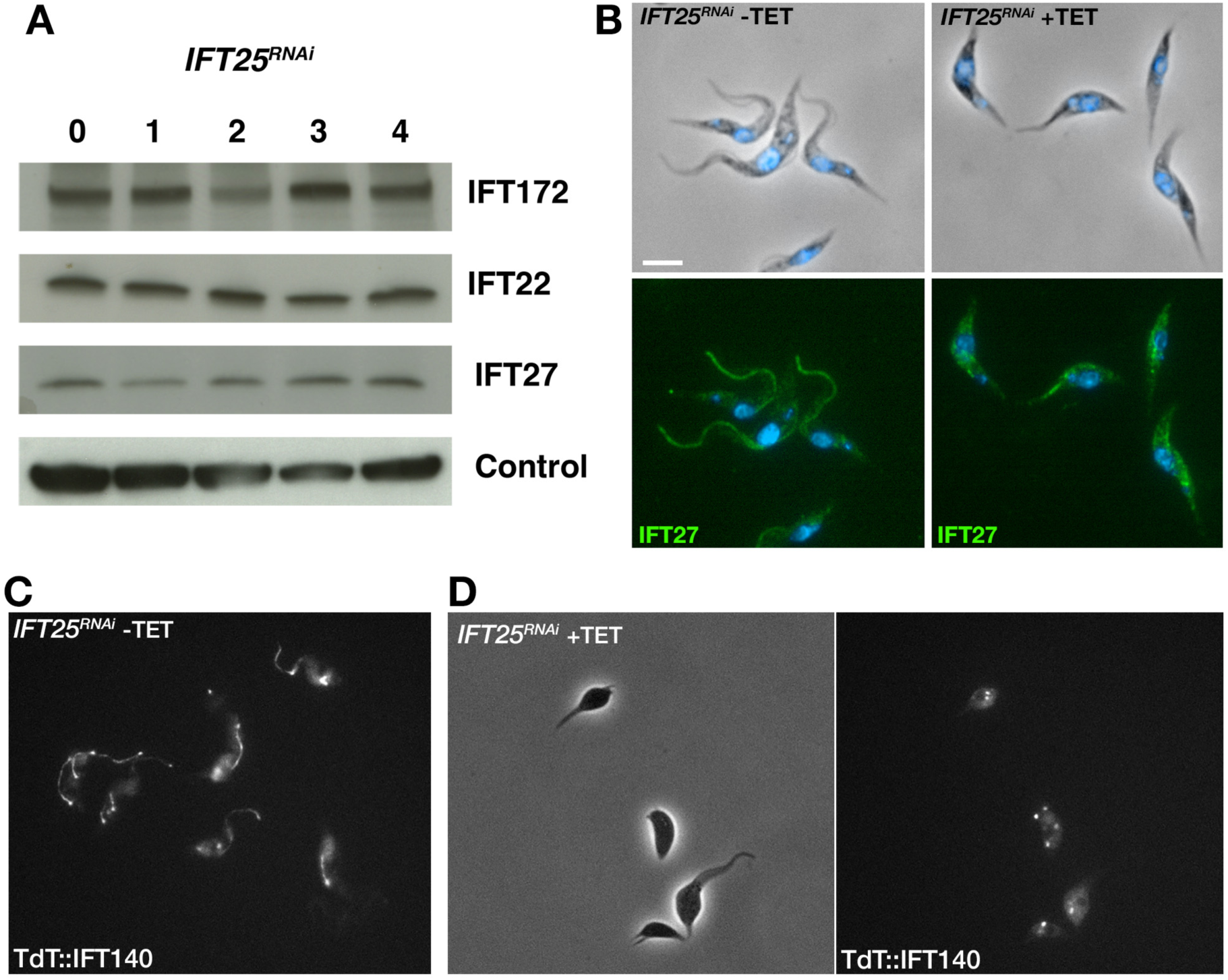
IFT27 and the IFT-A protein IFT140 fail to access the flagellum in the absence of IFT25. (A) Western blot using total protein extracts from the *IFT25^RNAi^* cell line induced for the indicated number of days. The membrane was blotted with the anti-IFT172 (top lane), anti-IFT22 (middle lane) and anti-IFT27 antibodies (bottom lane). The anti-PFR L13D6 antibody was used as loading control. (B) *IFT25^RNAi^* cells were non-induced (left) or induced for 3 days (right), labelled with the anti-IFT27 antibody and counterstained with DAPI. IFT27 does not accumulate in the short flagella but instead is dispersed in the cell body. Scale bar: 5µm. (C-D). *IFT25^RNAi^* cells were transformed to express the TdT::IFT140 from its endogenous locus and grown in non-induced conditions (C) or in the presence of tetracycline for 2 days to induce RNAi (D). Images are from Video S2 (C) or S3 (D). Sustained IFT trafficking of TdT::IFT140 is obvious in non-induced controls (C) whereas the fusion protein remains at the base of the flagella in induced cells.

## Discussion

Our functional analysis reveals that the trypanosome IFT25 is a *bona fide* IFT protein: (1) it is found within the matrix of the flagellum and concentrated at its base; (2) it traffics in both anterograde and retrograde directions at speed and frequency similar to those reported for other IFT proteins; (3) it is associated with other IFT-B proteins as shown by immuno-precipitation and bimolecular fluorescence complementation assays and (4) it is required for flagellum construction. Similarly to its partner IFT27/RABL4 (Huet et al., 2014) and to the other small G protein IFT22/RABL5 (Adhiambo et al., 2009), the knockdown of IFT25 restricts flagellum formation to short disorganised structures filled with electron-dense material that is positive for the IFT-B marker IFT172, indicative of a defect in retrograde transport. IFT27 did not accumulate in the short flagella of induced *IFT25^RNAi^* cells but instead was dispersed in the cytoplasm. This is agreement with *in vitro* and *in vivo* data in *Chlamydomonas* that showed tight interactions between IFT25 and IFT27 prior to their association to the IFT-B complex (Bhogaraju et al., 2011; Wang et al., 2009). The mislocalisation of the IFT-A marker IFT140 at the base of the short dilated flagella in the IFT25 knockdown cells suggests an uncoupling between IFT-B and IFT-A complexes that could explain the observed retrograde transport phenotype similar to *IFT27^RNAi^* (Huet et al., 2014).

This phenotype is quite similar to what was reported upon conditional deletion of IFT25 or IFT27 in male germ cells in mouse where spermatozoa possessed much shorter flagella that displayed numerous structural defects (Liu et al., 2017; Zhang et al., 2017). These include aberrant and disorganised axonemes that were too short, defects in associated extra-axonemal structures such as the outer dense fibres or the fibrous sheath and presence of vesicles in the lumen of the flagellum. Moreover, scanning electron microscopy revealed dilation of unknown molecular composition at the distal end, leading the authors to suggest a defect in retrograde IFT (Liu et al., 2017; Zhang et al., 2017). All these defects are quite similar to what is reported here for the knockdown of IFT25 in *T. brucei*, except that the nature of the accumulated material in aberrant spermatozoa has not been characterised in molecular composition. In both cases, these flagella were immotile.

In contrast to these results in spermatozoa and trypanosomes, IFT25 and IFT27 are not required for axoneme assembly in different mouse ciliated cells but are essential for hedgehog signalling (Keady et al., 2012; Yang et al., 2015), probably via trafficking of the BBSome (Eguether et al., 2014; Liew et al., 2014). Accordingly, a point mutation in the *IFT27* gene in humans resulted in Bardet-Biedl Syndrome (Aldahmesh et al., 2014). By contrast, a predicted biallelic loss of IFT27 led to a much more severe ciliopathy in a foetus (Quelin et al., 2018). In *Chlamydomonas*, IFT25 was shown to undergo IFT in the flagellum but inhibition of its expression using a microRNA did not interfere with flagellum formation, although defects in export of the BBSome from the flagella were observed (Dong et al., 2017).

We can propose at least three hypotheses as to why IFT25 could be essential for flagellum assembly in trypanosomes and mouse spermatozoa, and not in the other ciliated cells examined so far. First, the composition or organisation of IFT trains could be different between these cell types. Differences in train composition could be issued from variant mRNA resulting from different splicing, or to post-translational modifications of IFT proteins, to a modification in their stoichiometry or to the presence of additional proteins. In *Chlamydomonas*, IFT25 exhibits different phosphorylation patterns and only the phosphorylated version co-sediments with IFT27 on sucrose gradients (Richey and Qin, 2012). If the trains are different, one could envisage that the loss of IFT25/IFT27 has consequences on association with other IFT components. For example, the interaction between the IFT-B complex and the dynein cargo (Jordan et al., 2018) could be dependent of IFT25/IFT27 is some assemblies and not in others. Loss of the interaction would lead to a failure in importing dynein and therefore in a retrograde transport defect.

In a second case, IFT25/IFT27 could be required for the transport of specific cargoes that are expressed in only certain cell types. These cargoes could be members of the Hedgehog pathway in several mouse cells (Keady et al., 2012; Yang et al., 2015). Another potential cargo could be the BBSome that shows occasional but consistent association with IFT trains in *Chlamydomonas* (Lechtreck et al., 2009a). Of note, deletion of *BBS* genes in mouse does not have a visible impact on primary cilia in the kidney and on motile cilia in the trachea but leads to absence of the sperm flagellum (Mykytyn et al., 2004), reminiscent to what was reported for the deletion of IFT25 or IFT27. If the later two contribute to the trafficking of BBS proteins, one could envisage that the spermatozoa phenotype is due to BBS perturbation. In trypanosomes, BBS proteins are not found in the flagellum but are located at the flagellar pocket membrane and the endocytic network (Langousis et al., 2016). Knockout of *BBS* genes does not impact flagellum construction and parasites even infect mice normally (Langousis et al., 2016). Therefore, defects in BBS trafficking are unlikely to be responsible for the failure of assembling the flagellum. It should be noted that the difference in cargo loading could be quantitative rather than qualitative. The sperm flagellum is very long (100 µm in mouse (Gage, 1998)) and might require transportation of large quantities of cargoes that would not be necessary for shorter cilia. Possibly IFT25 and IFT27 favour the loading of larger quantities of cargoes. The trypanosome flagellum measures 20 µm (Robinson et al., 1995), which is fairly long too.

A third possibility is an intrinsic difference between the organelles themselves: trypanosome and spermatozoa flagella are characterised by the presence of extra-axonemal structures (Escalier, 2003; Portman and Gull, 2010) that are not found in the other cases discussed here. These structures could impact on the displacement of IFT trains or perhaps require specific adaptors for transport of their own components in the flagellum (Kierszenbaum, 2002). Alternatively, defects in their construction and accumulation of aberrant structures could interfere with proper IFT of tubulin and other axonemal components and subsequently impact on axoneme construction that would hence be a secondary phenotype. In *T. brucei*, absence of one of the two main PFR proteins leads to the accumulation of the other one at the distal tip of the flagellum (Bastin et al., 1999; Bastin et al., 1998), suggesting that PFR proteins are indeed transported. However, the axoneme looked normal and IFT trains were detected by transmission electron microscopy (Bastin et al., 1999). Major disruptions of the PFR have been reported upon RNAi knockdown of calmodulin (Ginger et al., 2013) or of the kinesin KIF9B (Demonchy et al., 2009). These flagella appear shorter than normal but this is less pronounced compared to the tiny flagella assembled in the *IFT25^RNAi^* or *IFT27^RNAi^* cells (Huet et al., 2014). Possibly the presence of extra-axonemal structures imposes constraints on IFT. In trypanosomes, electron microscopy analysis revealed that IFT trains are only encountered on two doublets (Bertiaux et al., 2018), whereas they are present on most of them in *Chlamydomonas* (Stepanek and Pigino, 2016). These two doublets are the one directly adjacent to the PFR (Sherwin and Gull, 1989). To our knowledge, IFT trains have not been visualised at the electron microscopy level in mammalian cells. The comparison of IFT trains between cilia of the trachea and the flagellum of growing spermatozoa could be promising. It should be reminded here that IFT proteins are only detected during construction of the flagellum and are absent of mature spermatozoa (San Agustin et al., 2015).

In conclusion, IFT25 and IFT27 are two IFT proteins that are associated to the IFT-B complex and participate to IFT. Their contribution to axoneme construction varies from one cell type to the other and so far they appear crucial for long organelles containing extra-axonemal structures. Nevertheless, they can play a role in other functions such as the hedgehog pathway in mouse. Their loss in multiple eukaryotic lineages (van Dam et al., 2013) remains intriguing. Investigation of their role in different organisms with different types of cilia will be necessary to have a more global view of the biological role of IFT25/IFT27 and to understand their loss is so many species.

## Materials and methods

### Cell lines and culture conditions

All procyclic *T. brucei* cell lines were derivatives of strain 427 and grown in SDM79 medium with hemin and 10% fetal calf serum. The 29-13 cell line expressing the T7 RNA polymerase and the tetracycline-repressor (Wirtz et al., 1999) has been described previously. For generation of the *IFT25^RNAi^* cell line, the full *IFT25* gene (Tb927.11.13300) consisting of 471 nucleotides was chemically synthesized by GeneCust Europe (Dudelange, Luxembourg) and cloned in the pZJM vector (Wang et al., 2000), allowing for tetracycline-inducible expression of dsRNA generating RNAi upon transfection in the 29-13 recipient cell line. The dsRNA is expressed from two tetracycline-inducible T7 promoters facing each other in the pZJM vector. The plasmid was linearized at the unique NotI site in the rDNA intergenic targeting region before transfection (Wang et al., 2000). The synthesis of the IFT25 coding sequence was conducted by GeneCust Europe and cloned into the pPCPFR vector (Adhiambo et al., 2009) to generate the pPCPFReGFPIFT25 plasmid allowing a GFP tagging at the N-terminal of IFT25. The resulting plasmid was linearized with NsiI, targeting integration in the intergenic region of *PFR2*. For bimolecular fluorescence complementation assays, two constructs were prepared for tagging IFT25 with the first part of YFP and IFT27 with the second part. If the two proteins interact closely, the YFP should be reconstituted and emit light (Kerppola, 2008). The first 465 nucleotides of *YFP* were synthesised in a fusion at the 5’ end of the first 498 nucleotides of the *IFT25* coding sequence and cloned in the p2675 vector (Kelly et al., 2007) for *in situ* insertion in the *IFT25* gene upon linearization with EcoRV and transfection in wild-type cells, resulting in the expression of the YFP^Nter^::IFT25 protein. The last 252 nucleotides of YFP, preceded by an ATG start codon, were synthesised fused to the sequence of the Ty1 tag (Bastin et al., 1996) and followed by the first 440 nucleotides of the *IFT27* coding sequence, resulting in the expression of the YFP^Cter^::IFT27 protein. The synthesised product was cloned in the p2845 vector (Kelly et al., 2007) and linearised with EcoRI to insert in the *IFT27* locus transfection in wild-type cells. Screening with a polyclonal anti-GFP antibody (Invitrogen) confirmed the presence of each fusion protein. The cell line expressing the YFP^Nter^::IFT25 protein was next transfected with the construct for expressing of the YFP^Cter^::IFT27 protein and resistant cells were screened using live fluorescence imaging. Trypanosomes were transfected with the plasmid constructs by Nucleofector technology (Lonza, Italy) (Burkard et al., 2007). Transfectants were grown in medium with the appropriate antibiotic concentration and clonal populations were obtained by limited dilution.

### Scanning and transmission electron microscopy

For transmission electron microscopy, cells were fixed in culture media overnight at 4°C with 2.5% glutaraldehyde in 0.1 M cacodylate buffer (pH 7.2) and then post-fixed in OsO4 (2%) in the same buffer. After serial dehydration, samples were embedded in Agar 100 (Agar Scientific, Ltd., United Kingdom) and left to polymerize at 60°C. Ultrathin sections (50–70 nm thick) were collected on Formvar-carbon-coated copper grids using a Leica EM UC6 ultra-microtome and stained with uranyl acetate and lead citrate. Observations were made on a Tecnai 10 electron microscope (FEI) and images were captured with a MegaView II camera and processed with AnalySIS and Adobe Photoshop CS4 (San Jose, CA). For scanning electron microscopy, samples were fixed overnight at 4°C in 2.5% glutaraldehyde as described previously (Absalon et al., 2007). Observations were made in a JEOL 7600F microscope.

### Immunofluorescence and live cell imaging

Cultured parasites were washed twice in SDM79 medium without serum and spread directly into poly-L-lysine coated slides. The slides were air-dried for 10 minutes, fixed in methanol at −20°C for 30 seconds and rehydrated for 10 min in PBS. For immunodetection, slides were incubated with primary antibodies diluted in PBS containing 0.1% (w/v) Bovine Serum Albumin (BSA) for 1 hour. Three washes of 10 min were performed and the secondary antibody diluted in PBS with 0.1% BSA was added to the slides. After an incubation of 45 min, slides were washed three times in PBS for 10 min and DAPI (2µg/µL) was added. Slides were mounted using ProLong antifade reagent (Invitrogen). Antibodies used were: mouse anti-IFT27 diluted 1/800 (Huet et al., 2014), mAb25 recognizing TbSAXO1, a protein found all along the trypanosome axoneme (Dacheux et al., 2012), the anti-IFT172 mouse monoclonal antibody (Absalon et al., 2008b) and a commercial rabbit polyclonal anti-GFP (Invitrogen). Subclass-specific secondary antibodies coupled to Alexa 488 and Cy3 (1/400; Jackson ImmunoResearch Laboratories, West Grove, PA) were used for double labelling. Sample observation was performed using a DMI4000 microscope (Leica) and images acquired with an ORCA-03G camera (Hamamatsu, Hamamastu City, Japan). Pictures were analysed using ImageJ 1.47g13 software (National Institutes of Health, Bethesda, MD) and images were merged and superimposed using Adobe Photoshop CC. For live video microscopy, cells were picked up from the culture, deposited on slides and covered by a coverslip, and observed directly with the DMI4000 microscope. Videos were acquired using an Evolve 512 EMCCD Camera (Photometrics, Tucson, AZ) and the Metamorph acquisition software (Molecular Probes, Sunnyvale, CA).

### Kymograph analyses

Trypanosomes were observed with the DMI4000 microscope and videos of IFT trafficking were recorded with a 250ms exposure per frame during 30s using an Evolve 512 EMCCD Camera (Photometrics, Tucson, AZ). Kymographs were extracted and analysed as described previously (Buisson et al., 2013; Chenouard et al., 2010).

### Western Blot

Cells were washed in PBS and boiled in Laemmli loading buffer before SDS-PAGE separation, loading 20 µg of total cell protein per lane. Proteins were transferred overnight at 25V at 4°C to polyvinylidene fluoride membranes, then blocked with 5% skimmed milk in PBS containing Tween 0.1% (PBST) and incubated with primary antibodies diluted in 1% milk and PBST. The anti-IFT27 serum was diluted 1/800, and the anti-IFT172 was diluted 1/500. To detect GFP, we used an anti-GFP antibody (Roche) diluted 1/500. As loading controls, antibodies against PFR proteins (L13D6)(Kohl et al., 1999) diluted 1/50 and anti-aldolase (a kind gift of Paul Michels, Brussels, Belgium) diluted 1/1000 were used. Three membrane washes were performed with PBST for 5 min then species-specific secondary antibodies coupled to horseradish peroxidase (GE Healthcare) were added at a 1/20 000 dilution in PBST containing 1% milk and incubated for 1 h. Final detection was carried out by using an enhanced chemo-luminescence kit and a high performance chemo-luminescence film according to manufacturer’s instructions (Amersham, Piscataway, NJ).

### Immunoprecipitation

50 mL of WT cells expressing GFP::IFT25 were grown at a final concentration of 1.10^7^ cells/ml. Cells were washed and incubated in IP_2_ buffer (0.8M NaCl, 20 mM Tris-HCl pH 8, 20mM EDTA, 2% Triton X100, 10µg/mL leupeptin and pepstatine A) for 30 min at room temperature. Soluble protein extract was centrifuged at 21,130*g* at 4°C for 30 min and supernatant was collected and incubated for 12h at 4°C with either 5µl of polyclonal rabbit anti-GFP (Invitrogen) or 5µl of anti-aldolase as control. Prior to the immune complex recovery, protein A-Sepharose beads (GE Healthcare, NJ) were washed in IP_1_ buffer (0.4M NaCl, 10 mM Tris-HCl pH 7,4, 10mM EDTA, 1% Triton X100, 1mg/mL ovalbumine). Immune complexes were recovered by incubation with pre-treated protein A-Sepharose beads for 15 min at 4°C. The beads then were washed three times with IP_1_ buffer, twice with IP_2_ buffer and once with IP_3_ buffer (0.4M NaCl, 10 mM Tris-HCl pH 7,4, 10mM EDTA). Interaction of the immune complex with the beads was disrupted by boiling in 50 µL of Laemmli loading buffer and analysed by SDS-PAGE followed by western blotting.

## Acknowledgements

We thank Monica Bettencourt-Dias and Esben Lorentzen for useful discussions. We thank the Ultrapole/Ultrastructural Bioimaging of the Institut Pasteur for providing access to their equipment.

This work is funded by ANR grants (11-BSV8-016 and 14-CE35-0009-01), by a French Government Investissement d’Avenir programme, Laboratoire d’Excellence “Integrative Biology of Emerging Infectious Diseases” (ANR-10-LABX-62-IBEID) and by La Fondation pour la Recherche Médicale (Equipe FRM DEQ20150734356). Diego Huet was funded by doctoral fellowships from the French Ministry for Research and from La Fondation pour la Recherche Médicale (FDT20120925362).

## Author contributions

D.H. and P.B. designed research; D.H., T.B. and S.P. performed research; D.H., T.B. and P.B. analysed data; D.H. and P.B. wrote the paper.

The authors declare no conflict of interest.

**Supplementary figure S1.**
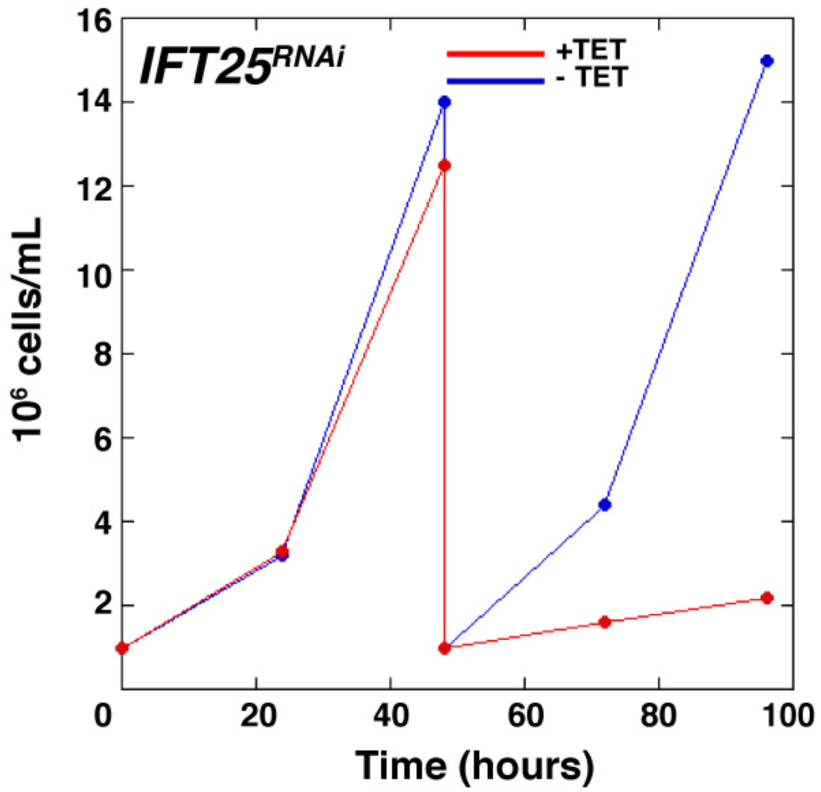
IFT25 is essential for growth in *T. brucei.* Growth curve of non-induced (blue) and induced (red) *IFT25^RNAi^* cells revealed very slow growth after two days in RNAi conditions, i.e. after the emergence of non-flagellated cells. Cells were diluted at day 2 of the experiment.

## Legends for supplementary videos

Video S1. Trypanosome expressing GFP::IFT25. IFT trafficking was recorded with a 250ms exposure per frame.

Video S2. Trypanosomes expressing YFPNter::IFT25 and YFPCter::IFT27. IFT trafficking was recorded with a 250ms exposure per frame.

Video S3. *IFT25^RNAi^* cell expressing TdT::IFT140 grown in the absence of tetracycline and hence without RNAi. Robust IFT trafficking is visible in all cells.

Video S4. *IFT25^RNAi^* cell expressing TdT::IFT140 grown in the presence of tetracycline and hence in RNAi conditions. No trafficking is observed and TdT::IFT140 appears concentrated in the basal body area.

